# Pathogenicity and transmissibility of bovine-derived HPAI H5N1 B3.13 virus in pigs

**DOI:** 10.1101/2025.03.04.641414

**Authors:** Taeyong Kwon, Jessie D. Trujillo, Mariano Carossino, Heather M. Machkovech, Konner Cool, Eu Lim Lyoo, Gagandeep Singh, Sujan Kafle, Shanmugasundaram Elango, Govindsamy Vediyappan, Wanting Wei, Nicholas Minor, Franco S. Matias-Ferreyra, Igor Morozov, Natasha N. Gaudreault, Udeni B. R. Balasuriya, Lisa Hensley, Diego G. Diel, Wenjun Ma, Thomas C. Friedrich, Juergen A. Richt

**Author notes:** Corresponding author: Dr. Juergen A. Richt.

## Abstract

Since the first emergence of highly pathogenic avian influenza (HPAI) H5N1 viruses in dairy cattle, the virus has continued to spread, reaching at least 17 states and at least 970 dairy herds in the United States. Subsequently, spillovers of the virus from dairy cattle to humans have been reported. Pigs are an important reservoir in influenza ecology because they serve as a mixing vessel in which novel reassortant viruses with pandemic potential can be generated. Here, we show that oro-respiratory infection of pigs resulted in productive replication of a bovine-derived HPAI H5N1 B3.13 virus. Infectious virus was mainly identified in the lower respiratory tract of principal infected pigs, and sero-conversion was observed in most of the principal pigs at later time points. In one animal, we detected the emergence of a mutation in hemagglutinin (HA) previously associated with increased affinity for “mammalian-type” α2,6-linked sialic acid receptors, but this mutation did not reach consensus levels. Sentinel contact pigs remained sero-negative throughout the study, indicating lack of transmission. The results support that pigs are susceptible to a bovine-derived HPAI H5N1 B3.13 virus, but this virus did not replicate as robustly in pigs as mink-derived HPAI H5N1 and swine-adapted influenza viruses.

## Introduction

Highly pathogenic avian influenza (HPAI) viruses have multibasic amino acids in the cleavage site of the HA protein that are recognized by ubiquitous proteases, resulting in systemic disease in poultry [1]. Since the first emergence of A/Goose/Guangdong/1/1996 (Gs/Gd) in 1996, the Gs/Gd lineage of HPAI viruses have sporadically infected humans and caused a 40 to 50% case fatality rate in laboratory-confirmed humans [2]. After continuous evolution, the HA genes of the clade 2.3.4.4b became dominant in HPAI H5 viruses, and spread across the globe via migratory birds, causing HPAI outbreaks in Asia, Europe and North America since 2014 [3,4]. More recently, spillover events of HPAI H5 clade 2.3.4.4b to a variety of mammalian species, especially carnivores and aquatic mammals, have been identified [5]. In early 2024, the first case of HPAI infection in cattle was reported in Texas, United States. Infected dairy cattle exhibited fever, loss of appetite, reduced milk production and poor milk quality [6]. The causative agent was identified as HPAI H5N1 genotype B3.13 virus that was likely the result of a reassortment event between a HPAI H5N1 B3.6 ancestor and North American LPAI viruses [6–8]. On January 2025, a new genotype D1.1 was first detected in bulk milk samples, and trace investigation confirmed HPAI outbreaks caused by D1.1 virus in Nevada dairy farms [9]. As of 2/28/2025, the virus readily spread across the country, affecting 976 dairy herds in 17 states [10]. Moreover, 68 human cases have been reported and the infected humans showed in general mild disease with a single fatal case so far [11]. Most cases were linked to direct exposure to infected cattle (n=41) or poultry (n=24). Experimental studies were immediately performed in cattle to determine the pathogenicity and transmissibility of the newly emerged B3.13 viruses. Experimental infection of lactating cattle via intra-mammary route recapitulated the disease observed in the field, whereas calves infected through intra-nasal and oral routes showed only mild disease with limited virus replication and no transmission to contact sentinels. These findings support the hypothesis that natural infection is established via the intra-mammary route, potentially through the milking process on dairy farms [12,13].

Pigs are natural hosts for H1N1, H1N2, and H3N2 subtypes of influenza A viruses (IAVs) and these viruses readily transmit in pig populations [14]. Sporadic H5N1 infection in pigs has been documented [15,16]. Pigs express sialic acid receptors for both mammalian and avian IAVs in their respiratory epithelium, enabling some avian IAVs to infect pigs [17]. The introduction of avian and human IAVs into swine populations and subsequent reassortment facilitate the expansion of genetic diversity of swine IAVs. Modern agricultural systems and swine exhibitions facilitate spillover events of IAV from swine to humans or vice versa, forming intertwined clusters in phylogenetic trees [18]. Thus, pigs and humans frequently exchange IAV lineages, while pigs are also more susceptible than humans to avian-origin IAVs. Because of this, pigs are thought to serve as mixing vessels in which novel viruses with pandemic potential can be generated [19]. The important role of pigs as the mixing vessel is exampled by the 2009 H1N1 pandemic, where the virus underwent reassortment in pigs and acquired novel features and antigenicity to evade the pre-existing immunity in humans [20].

Although the current agricultural system enables inter-species transmission of IAV to pigs, the potential of the bovine-derived HPAI H5N1 clade 2.3.4.4b viruses for infection and transmission in pigs remains unclear. Notably, the US first case of H5N1-infected pig was identified in a backyard farm in Oregon, suggesting the expansion of the host range of the contemporary HPAI H5N1 viruses to agricultural animals [21]. Therefore, we aimed to evaluate the pathogenicity and transmissibility of the bovine-derived HPAI H5N1 clade 2.3.4.4b genotype B3.13 virus in pigs.

## Materials and methods

### Viruses and cells

Madin-Darby canine kidney (MDCK) cells were maintained in Dulbecco’s Modified Eagle Medium (DMEM; Corning, Manassas, VA, USA) supplemented with 5% fetal bovine serum (FBS; R&D systems, Flower Branch, GA, USA) and a 1% antibiotic-antimycotic solution (Gibco, Grand Island, NY, USA). The bovine-derived HPAI H5N1 B3.13 virus, A/Cattle/Texas/063224-24-1/2024, was propagated on bovine uterine epithelial cells (CAL-1; In-house) for 3 passages at Cornell University [6]. At KSU, the virus was propagated on MDCK cells for 1 passage in influenza infection media (DMEM supplemented 0.3% bovine serum albumin, 1% antibiotics-antimycotic solution, 1% non-essential vitamin and 1 µg/mL of TPCK-treated trypsin).

### Animal study

The animal study and experiments were approved and performed under the Kansas State University (KSU) Institutional Biosafety Committee (IBC, Protocol #1545) and the Institutional Animal Care and Use Committee (IACUC, Protocol #4778) in compliance with the Animal Welfare Act. All animal and laboratory work were performed in a biosafety level-3+ laboratory and BSL-3Ag facility in the Biosecurity Research Institute at KSU in Manhattan, KS, USA except for inactivated samples at BSL-2. Twelve commercial 4-week-old Yorkshire crossbred piglets were obtained from a herd with high health status. Pigs were randomly assigned into principal-infected (n=9) and sentinel (n=3) groups. After acclimation, principals were challenged with 5 × 10^6^ TCID_50_ per pig via intra-tracheal (2 mL), intranasal (1 mL), and oral (1 mL) routes. At 2 day post-challenge (DPC), three sentinels were commingled with principals in a single pen to allow direct contact. Nasal swabs were collected daily until the end of study and oropharyngeal swabs were collected at 1, 3, 5, 7, 10, and 14 DPC. The swabs were resuspended in 2 mL of DMEM supplemented penicillin and streptomycin, filtered using a 0.45µm syringe filter and stored – 80 °C until further analysis. Serum was collected for evaluating antibody responses. Six pigs were randomly selected and humanly euthanized at 3 (n=3, pig ID: #50, #52, and #54) and 5 DPC (n=3, #51, #53, and #61), and the remaining three principals (#57, #59, and #60) and three sentinels (#55, #56, and #58) were humanly euthanized at 14 DPC. At necropsy, bronchoalveolar lavage fluid (BALF) and tissue samples were collected for virological and pathological evaluation.

### Virus titration

Nasal swab, oropharyngeal swab, BALF and 10% tissue homogenates (w/v) were subjected to virus titration. Clarified or filtered samples were serially diluted in influenza infection media. After washing with 1X PBS, the confluent MDCK cells were inoculated with serially diluted samples. On day 2, the cells were fixed with 100% cold methanol for immunofluorescence assay (IFA) and incubated with the primary antibody (mouse monoclonal antibody HB65, specific for IAV nucleoprotein), and then with anti-mouse IgG conjugated with Alexa Fluor 488 (Invitrogen, Carlsbad, CA, USA). The virus-positive cells were visualized on an EVOS system. The virus titer was determined using the Reed-Muench method.

### RNA extraction and Influenza A Matrix-specific RT-qPCR

To determine viral RNA shedding and distribution, viral RNA was extracted using a magnetic bead-based automated extraction system (Taco Mini and total nucleic acid extraction kit, Gene Reach, USA) as previously reported [22]. Briefly, nasal swabs, BALF samples and 10% tissue homogenates (DMEM) or formalin fixed tissue homogenates in AL lysis buffer and digested with proteinase K were mixed with an equal volume of the RLT buffer (Qiagen, Germantown, MD, USA) and incubated for 20 minutes at room temperature. Two hundred microliters of the RLT lysate and 200 µL isopropanol were added to the wells containing magnetic beads, followed by 4 sequential washes prior to elution of nucleic acids from the magnetic beads using automated extraction machines (Gene reach Taco Mini or Qiagen Biosprint). RT-qPCR was performed on the Biorad CFX or Opus using a modified version of the NAHLN FLU A matrix detection assay utilizing the qScript XLT 1-Step RT-qPCR ToughMix (QuantaBio, Beverly, MA, USA) [22,23]. RT-qPCR was performed in duplicates with a Ct cut-off value of 38. Any samples with Ct>38 or those with positive results on a single replicate were considered suspect positive.

### Gross pathology and histopathology

Macroscopic lung lesions were scored, and the percentage of affected pulmonary parenchyma was calculated as previously described [22]. Tissue samples were fixed in 10% neutral buffered formalin, trimmed, processed and stained with hematoxylin and eosin (H&E) following standard procedures. Microscopic pathology was evaluated and scored based on microanatomy of the lung [22]. A minimum of 6 (sentinel animals) and a maximum of 16 (principal-infected animals) lung sections representing all lobes were scored from 0 to 4 (0 = absent, 1 = minimal, 2 = mild, 3 = moderate, and 4 = severe) for six separate criteria typically associated with IAV infection in pigs. Scoring categories include (i) epithelial necrosis or attenuation; (ii) airway exudate; (iii) percentage of airways with inflammation; (iv) peribronchiolar and perivascular lymphocytic inflammation; (v) alveolar exudate; (vi) alveolar septal inflammation. For scoring the percentage of airways with inflammation (iii): 0% = 0, up to 10% involvement = 1, 10– 39% involvement = 2, 40–69% involvement = 3 and, greater than 70% involvement = 4. Additional tissues from the respiratory tract and lymphoid system were evaluated.

### Influenza A virus-specific immunohistochemistry (IHC)

Immunohistochemistry (IHC) for detection of IAV H5N1 nucleoprotein (NP) antigen was performed on the automated BOND RXm platform using the Polymer Refine Red Detection kit (Leica Biosystems, Buffalo Grove, IL) as described previously [22]. The primary antibody (rabbit monoclonal anti-Influenza A virus NP [Cell Signaling Technology, #99797/F8L6X]) was diluted 1:1,200 in Primary Antibody diluent (Leica Biosystems) and incubated for 30 min at ambient temperature. Lung sections from a goose naturally infected with a HPAI H5N1 strain submitted to the Louisiana Animal Disease Diagnostic Laboratory (LADDL) were used as positive assay control.

### Serology

The virus neutralization test was performed according to a previously established protocol [24]. Briefly, serum samples were heat-inactivated at 56 °C for 30 minutes. A total of 50 µL of a two-fold diluted serum sample was mixed with the equal volume of 100 TCID_50_ of the virus and incubated at 37 °C for 1 hour. After incubation, the mixture was transferred to pre-washed MDCK cells. On day 2, IFA was performed as mentioned above, in order to determine 50% inhibition of virus growth. In addition, commercial NP-based enzyme-linked immunosorbent assay (ELISA) kits, including three competition ELISA kits and one indirect ELISA kit, were used to determine the serological response according to the manufacturers’ instruction: (1) AsurDx™ Influenza A virus (IAV) Antibodies cELISA Test Kit, BioStone, USA, (2) ID Screen® Influenza A Nucleoprotein Swine Indirect, Innovative Diagnostics, France, (3) IDEXX Swine Influenza Virus Ab Test, IDEXX, USA and (4) ID Screen® Influenza A Antibody Competition Multi-Species, Innovative Diagnostics, France.

### Complementary DNA (cDNA) generation and PCR

We performed reverse transcription, PCR amplification, and library prep as described in the publicly available protocol (https://dx.doi.org/10.17504/protocols.io.kqdg322kpv25/v1). We performed sequencing in two technical replicates, using the same input RNA. We used reagents from the QIAseq DIRECT SARS-CoV-2 Kit A and QIAseq DIRECT SARS-CoV-2 Enhancer (Qiagen). For each sample, we first performed first-strand cDNA synthesis. We assembled a reaction master mix on ice with RP Primer (1 µL), Multimodal RT Buffer (4 µL), RNase Inhibitor (1 µL), nuclease-free water (8 µL), and EZ Reverse Transcriptase (1 µL). We added 5 µL of sample RNA to this reaction. The thermal cycling conditions were 25°C for 10 minutes, 42°C for 10 minutes, 85°C for 5 minutes, followed by a hold at 4°C. At this point we also added a no-template control sample consisting of nuclease-free water.

Next, cDNA was amplified using a custom primer pool that generates overlapping amplicons of ∼250bp. Primer pools was designed as previously described [25]. Briefly, PrimalScheme [26] was used to generate primers from a set of clade 2.3.4.4b, genotype B3.13 H5N1 sequences from the U.S. dairy cattle outbreak (https://github.com/andersen-lab/avian-infuenza). Because sequences including full-length noncoding regions (NCRs) were not available for clade B3.13 viruses, we manually designed primers targeting the 5’ and 3’ termini of each segment. To do this, we inferred likely sequences for the full NCRs of each gene segment based on publicly available sequences of clade 2.3.4.4b H5N1 viruses isolated from North American birds and mammals (https://github.com/moncla-lab/h5-sequencing-protocol-dev/blob/main/RT_and_PCR_protocol.md). Primers targeting the 5’ and 3’ termini of each segment also included short overhangs not present in the target sequence to enhance amplification efficiency.

We used first-strand cDNA from each sample as templated for two independent PCR reactions (one for each primer pool), which we set up using reagents from QIAseq DIRECT SARS-CoV-2 Kit A and QIAseq DIRECT SARS-CoV-2 Enhancer. For each PCR reaction we assembled a reaction master mix using 10 µM primer pool (4 µL), 5x UPCR buffer (5 µL), QN Taq Polymerase (1 µL) and nuclease-free water (7 µL). Eight microliters of cDNA were added to each reaction. The cycling conditions were 98°C for 2 minutes, then 4 cycles of 98°C for 20 seconds and 63°C for 5 minutes, followed by 29 cycles of 98°C for 20 seconds and 63°C for 3 minutes with a final hold at 4°C. A Qubit fluorometer (ThermoFisher) was used to quantify the double-stranded DNA after the reaction.

### Oxford Nanopore library prep and sequencing

We prepared libraries according to the protocol for the Oxford Nanopore Technology Native Barcoding Kit V14 (Ligation Sequencing Amplicons -Native Barcoding Kit 96 V14 (SQK-NBD114.96), 2022). The library was sequenced on a minION mk1C sequencer using an R10 flow cell.

### Processing of the raw sequence data, mapping, variant calling and variant annotation

A bespoke Nextflow pipeline was used to perform Oxford Nanopore base-calling, primer-trimming, and variant-calling (https://github.com/nrminor/oneroof) [27]. The pipeline reproducibly supplies software dependencies via Docker containers, Apptainer containers, or local conda environments built with the Pixi package manager. Base-calling is performed with the Oxford Nanopore Dorado base-caller v0.7.1. Reads are adapter- and primer-trimmed using CutAdapt v4.8 [28]. For the results presented here, we required a minimum read length of 150 bases and a maximum read length of 500 bases, along with a minimum average read quality of 10. Reads are aligned to the reference genome using minimap2 v2.28, allowing up to 2 primer mismatches between a given primer and the reference sequence [29]. The reference genome used for this piepline is A/Bovine/Texas/24-029328-01/2024 (H5N1), a sequence isolated from an infected cow’s milk early in the outbreak [30]. For each gene segment, we manually added the inferred NCR sequences, as well as the synthetic “overhang” sequences present in each primer, to the reference sequence to aid in read mapping. iVar v1.4.2 [31] was used to call variants and snpEff v5.2 [32] was used for variant-effect annotation. Variants were called using a minimum required depth of coverage of 20 reads and a minimum variant frequency of 0.1. Consensus sequences were called using samtools v1.5 [33], with a minimum depth of coverage of 20 and a minimum variant frequency of 0.5.

Additional utilities in SeqKit2 v2.8.2 [34], rasusa v2.0.0 [35], vsearch v2.28.1 [36] are used throughout the pipeline, alongside other tools. A full list of the pipeline’s dependencies can be found in the GitHub repository’s Python project configuration file, “pyproject.toml”.

### iSNV quality control

VCF files from the Nextflow pipeline were cleaned using a custom R script (R version 4.4.2) [37]. Samples with negative viral titers were removed from downstream analysis. Variants were filtered to include only nonsynonymous and synonymous variants. iSNVs were required to be at a frequency of 0.1 and have a depth of coverage of at least 20 reads. iSNVs were required to be present in both technical replicates with ≥10% frequency based on Grubaugh et al [38]. iSNVs were compared to the GISAID inoculum sequence (accession number EPI_ISL_19155861) [6].

## Results

In order to determine the pathogenicity and transmissibility of the H5N1 bovine isolate in pigs, animals were challenged with a total of 5 × 10^6^ TCID_50_ intra-orally, intra-nasally and intra-tracheally under experimental settings, and sentinels were co-housed with principals at 2 DPC to allow direct contact (Fig 1A). All principal-infected (n=9) and sentinel (n=3) pigs were active and healthy without apparent clinical signs throughout the study (data not shown). Infectious virus was isolated from nasal swabs of two principal-infected pigs: #50 with 3.16 × 10^2^ TCID_50_/mL at 2 DPC and #61 with 2.15 × 10^4^ TCID_50_/mL at 4 DPC and 3.16 × 10^3^ TCID_50_/mL at 5 DPC (Figure 1B). Oropharyngeal swabs of three pigs were found virus-positive at 1 DPC (#52, #59, and #61) and 3 DPC (#52) but the titers were at the limit of detection (Figure 1C). RT-qPCR showed that 7 of 9 principals were positive for viral RNA on at least one time point from 1 to 5 DPC in either the nasal or oropharyngeal swabs, suggesting low intermittent shedding (Supplementary figures 1A and 1B). Low levels of infectious virus were isolated from BALF samples of two pigs at 3 DPC (#50 and #52) and other two at 5 DPC (#51 and #61) (Figure 1D). The BALF samples of the virus-negative pigs (#54 at 3 DPC and #53 at 5 DPC) were positive and suspect-positive in RT-qPCR (Supplementary figures 1C and 1D). Low levels of infectious virus was also isolated from lungs of principal-infected pigs: 4.64 × 10^1^ TCID_50_/mL for #50 and #52 at 3 DPC and 1x10^2^ and 4.64 × 10^1^ TCID_50_/mL for #53 and #61 at 5 DPC. One trachea sample (#61) was virus-positive with a titer of 4.64 × 10^1^ TCID_50_/mL at 5 DPC. RT-qPCR data demonstrated that viral RNA was present in multiple respiratory tissues and lymph nodes on 3 and 5 DPC, and in one animal in the central nervous system on 5 DPC (Supplementary figures 1C and 1D). No infectious virus was isolated from principals at 14 DPC. All swab, BALF, and tissue samples collected from sentinels were negative for virus isolation but a nasal swab and an oral swab of #58 at 4 DPC and 7 DPC, respectively, were suspect positive in RT-qPCR.

**Figure 1.**
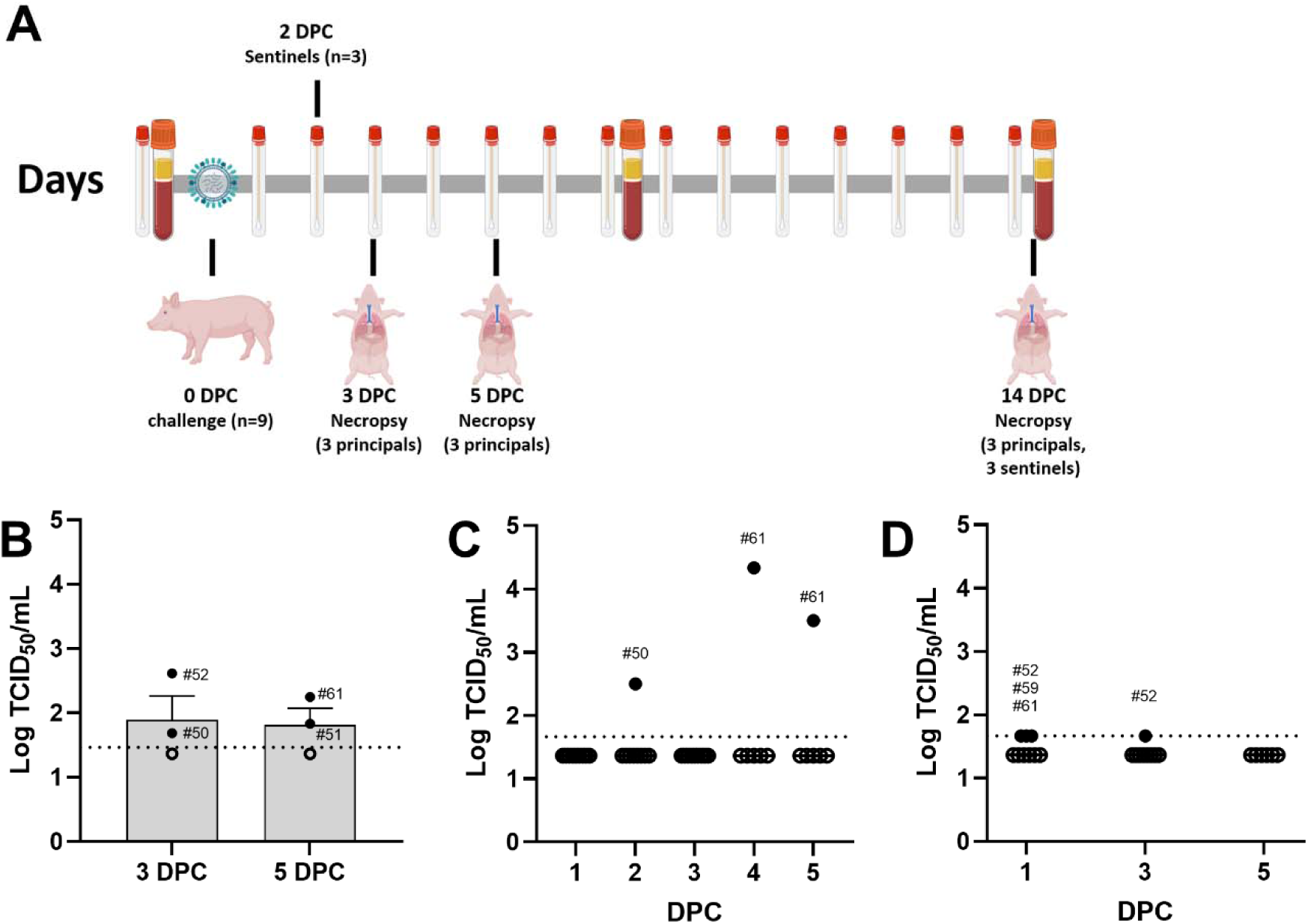
Study design and virus replication in pigs infected with a bovine-derived HPAI H5N1 B3.13 virus. (A) nine principals were challenged with 5 × 10^6^ TCID_50_ per pig via intra-tracheal (2 mL), intranasal (1 mL), and oral (1 mL) routes. Three sentinels were commingled with principals in a single pen at 2 DPC. Six pigs were randomly selected and humanly euthanized at 3 (n=3, pig ID: #50, #52, and #54) and 5 DPC (n=3, #51, #53, and #61), and the remaining three principals (#57, #59, and #60) and three sentinels (#55, #56, and #58) were humanly euthanized at 14 DPC. Nasal swab, oropharyngeal swab, and serum were collected throughout the study. Nasal swab (B), and oropharyngeal swab (C), and bronchoalveolar lavage fluid or BALF (D). Virus titration was performed on MDCK cells and virus-infected cells were visualized by immunofluorescence assay. The positive samples were shown as closed circles with pig ID; open circles represent below a limit of detection. The dash line is the LOD of assay (3.29 × 10^1^ TCID_50_/mL for BALF and 4.64 × 10^1^ TCID_50_/mL for swabs).

We next characterized pathological changes in principal-infected pigs at the time of necropsy. Macroscopically, pulmonary lesions in principal-infected pigs (6/6) were characterized by sporadic dark red and depressed lobules representing regions of lobular atelectasis typical of influenza virus A infections primarily at 3 DPC and less frequently at 5 DPC (Figure 2A and supplementary figure 2). Microscopic pulmonary lesions (Figures 2B and 3) consisted of mild to moderate multifocal non-suppurative interstitial pneumonia with rare necrotizing bronchitis/bronchiolitis affecting multiple bronchopulmonary segments. These alterations were mostly comprised of peribronchiolar and perivascular histiocytic and lymphocytic inflammation, extending to and expanding alveolar septa resulting in multifocal areas of consolidation (Figure 3A). In the affected bronchopulmonary segments, segmental or individual cellular degeneration and necrosis of the bronchiolar epithelium were noted at 3 DPC (figure 3A and lower right inset) and 5 DPC (Figure 3E upper right inset) in all principal-infected pigs. Intraluminal cell debris and neutrophils were rare. Affected bronchopulmonary segments were rarely accompanied by necrotizing alveolitis (Figure 3A lower left inset), collapse of the alveolar spaces and consolidation of the lobule at 3 DPC and 5 DPC (Figure 3E lower right inset). Areas with bronchiolar and alveolar microscopic changes were also characterized by intracytoplasmic and/or intranuclear viral antigen in bronchial/bronchiolar epithelial cells, pneumocytes and macrophages (3/3 principal-infected pigs at 3DPC; Figures 3B and 3C). At 5 DPC, histologic changes were similar to those occurring at 3DPC in all principal-infected pigs (3/3), with nearly absent bronchial and bronchiolar epithelial cell necrosis (Figures 3E and 3F). Viral antigen was most abundant at 3 DPC and identified in all principal-infected animals (Figures 3B and 3C) with immunolabeling of epithelia (alveolar and bronchiolar). By 5 DPC, viral antigen was present in all principal-infected animals but mostly limited to areas of alveolar/interstitial inflammation (Figure 3G), except in pig #51 and #61 where infrequent bronchiolar epithelial cells were sporadically immunolabeled. At 14 DPC, pulmonary changes were only at the level of the interstitium with multifocal to coalescing areas of mild to rarely moderate, non-suppurative interstitial pneumonia (3/3 principal-infected pigs). Alveolar septa were thickened by infiltrates of lymphocytes and macrophages (Figure 3H and inset) that also expanded interlobular septa (Figures 3H and 3I). Sporadic epithelial cells in bronchioles and alveoli as well as septal macrophages contained viral antigen (Figure 3J and 2/3 principal-infected pigs). Rare individual nasal (3 DPC #54), tracheal and bronchial (3 DPC# 50 and #54, 5 DPC #51 and #61) epithelial cells expressing viral antigen were noted. NP-specific IHC detected individual cellular staining in one or more of the tracheobronchial lymph nodes in all principal-infected pigs (Supplementary figure 3). Lower number of antigen-positive cells and weaker staining intensity were observed at 14 DPC, when compared to earlier time points. Rare scattered antigen-positive cells were also detected in the gastro-hepatic lymph nodes from #52 at 3 DPC and #51 at 5 DPC.

**Figure 2.**
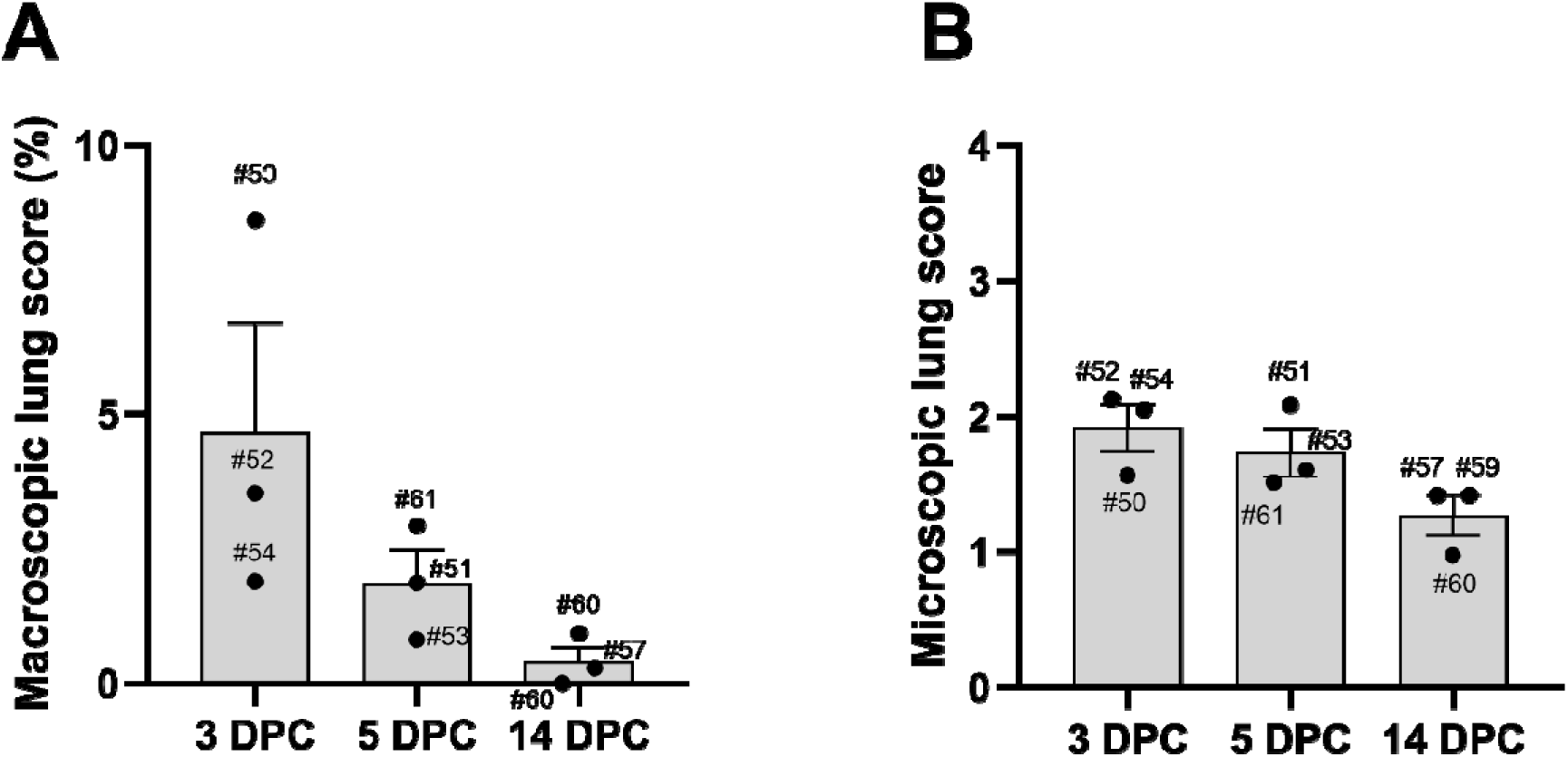
Macroscopic (A) and microscopic (B) lung scores in pigs infected with a bovine-derived HPAI H5N1 B3.13 virus. Macroscopic lung scores were calculated based on the percentage of the surface area showing lobular atelectasis and hyperemia. Microscopic scores were calculated by the average scores of six different criteria.

**Figure 3:**
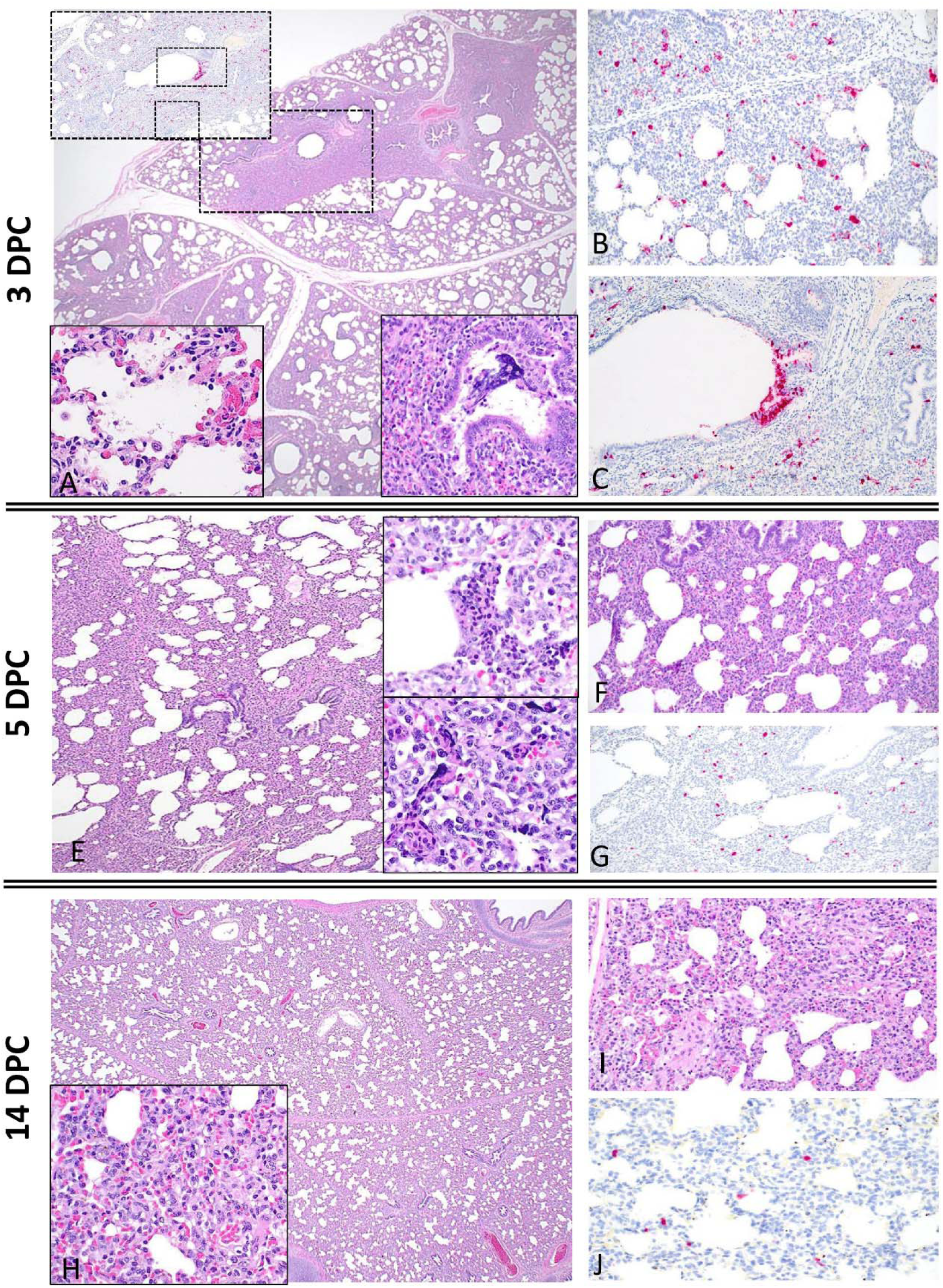
Histological features in lungs of pigs infected with a bovine-derived HPAI H5N1 B3.13 virus. Histologically, pulmonary lesions were characterized by multifocal to coalescing interstitial pneumonia with limited necrotizing bronchiolitis. At 3 DPC (A and lower right inset), multifocal bronchopulmonary segments were characterized by infrequent bronchiolar epithelial cell degeneration and necrosis (lower right inset) and associated areas of pulmonary parenchyma with collapsed alveolar spaces (atelectasis) and expanded alveolar septa by lymphocytes and histiocytes (A) and alveolar epithelial necrosis (A-lower right inset). Influenza A virus nucleoprotein intracytoplasmic antigen was detected within pneumocytes and alveolar macrophages (B) and segmentally or individually within bronchiolar epithelial cells (C). Infrequent yet similar epithelial necrosis was noted at 5 DPC (E, upper and lower insets), which is accompanied by multifocal to coalescing interstitial pneumonia characterized by alveolar septal thickening and cellular debris filling alveolar spaces (E and F) (F). Influenza A virus nucleoprotein antigen was much less abundant and mostly in cells lining and within alveoli (G). At 14 DPC, there are multifocal to coalescing regions areas of mild to moderate interstitial pneumonia. Alveolar septa are thickened by infiltrates of lymphocytes and macrophages and there is multifocal thickening of lobular septa with non-suppurative inflammation (H asterisk and I). Sporadic epithelial cells in bronchioles and alveoli as well as septal macrophages contained viral antigen (J). Magnifications of images: 20× for A, E and H, 100× for B, C, F, G, I and J. Magnifications of inserts: 400× for A, E and H and 100× for lower right (A).

There were no detectable neutralizing antibodies in the three principals at 7 DPC. Two principal pigs, #59 and #60, were sero-positive with 1:10 VN_50%_ titer at 10 DPC (Figure 4A). At 14 DPC, neutralizing antibodies were undetectable in serum of #60, but the other two principal pigs (#57 and #59) had detectable neutralizing antibodies against H5N1 viruses. Four different ELISA kits, including three NP-based competition ELISA kits and one NP-based indirect ELISA kit, supported clear seroconversion of principal pigs #57 and #59 at 14 DPC but not #60 (Figure 4B and supplementary figure 4). There was no antibody response in any of the sentinel pigs. Collectively, experimental infection led to low levels of antibody responses (#59 and #60) at 10 DPC and detectable neutralizing and NP-specific antibody responses in two (#57 and #59) of three principal-infected pigs at 14 DPC.

**Figure 4.**
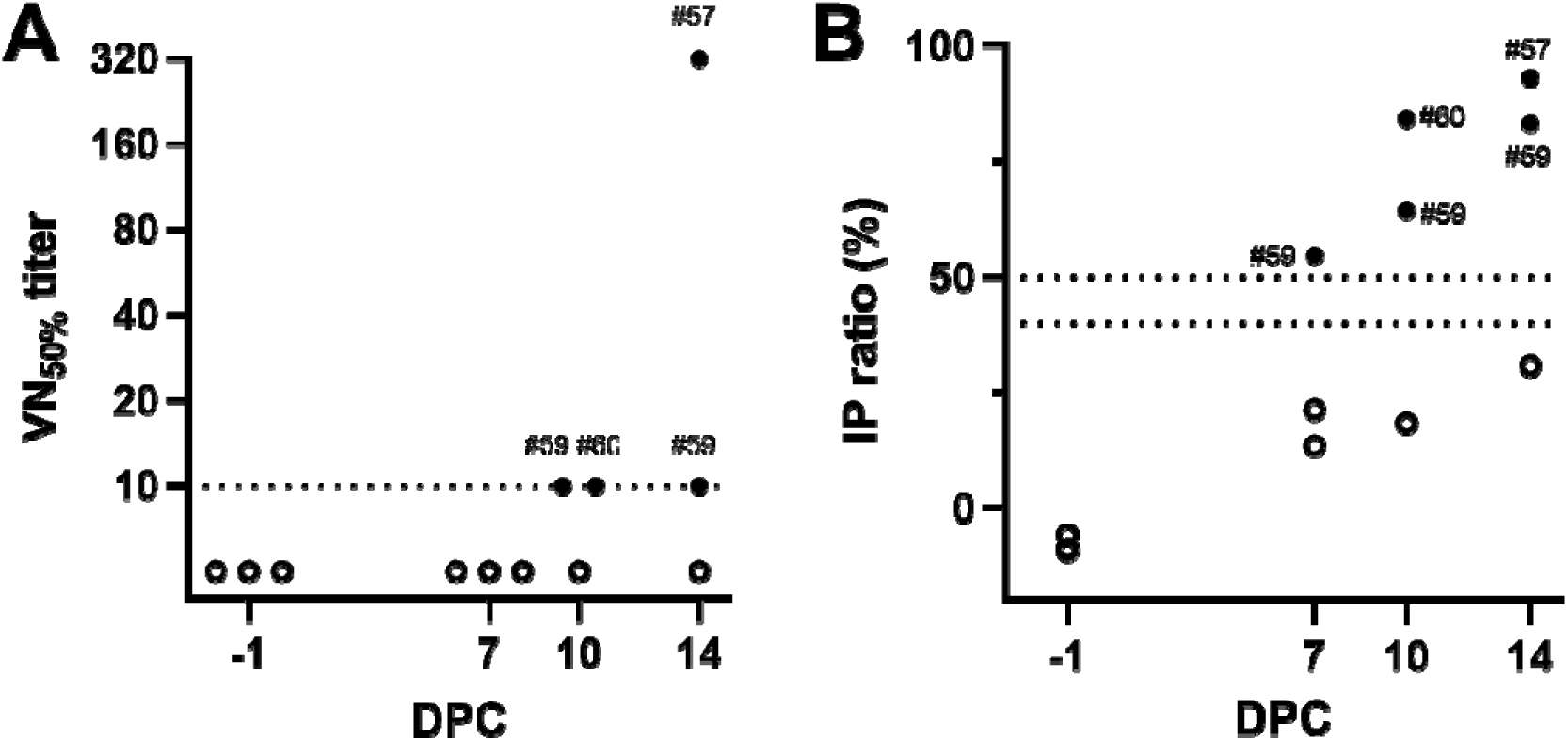
Antibody responses in pigs infected with a bovine-derived HPAI H5N1 B3.13 virus. Virus neutralization (VN) test (A) and commercial NP-based competition ELISA (B). (A) The virus neutralization test was performed with the challenge strain on MDCK cells to determine the 50% inhibition of virus growth. (B) Commercial NP-based competition ELISA (BioStone, USA) was used to determine serological responses. The positive samples were shown as closed circles with pig ID: open circles represent negative. The dash line is the initial dilution for the VN test (1:10) and the cut-off value of ELISA, <40%: negative, 40-50%: doubtful, >50% positive.

Finally, we asked whether selection might favor the accumulation of additional mammalian-adapting mutations in bovine-derived H5N1 viruses replicating in pigs. We generated viral genome sequences from all samples with sufficient vRNA in technical replicates using a previously described tiled amplicon approach and Oxford Nanopore sequencing (Supplementary figure 5). These sequences were analyzed using a custom pipeline that employs an early cattle isolate, A/Bovine/Texas/24-029328-01/2024, as a standard reference sequence. Note that this reference sequence is slightly different from the isolate used in our inoculations: the original isolate and the virus stock preparation used to inoculate our animals revealed6 mutations relative to this reference sequence. These mutations were maintained at the consensus level in all the specimens from infected pigs, with the exception of one synonymous PB2-A525G (cDNA position), which declined over time below consensus level in some pigs. In addition to these mutations, a total of 11 synonymous and 6 nonsynonymous *de-novo* consensus-level mutations emerged in infected pigs. Five of the 6 *de-novo* nonsynonymous mutations occurred in pig #61, which had the highest viral titers of all the animals (Figure 5). Among these 5 nonsynonymous mutations is an isoleucine-to-methionine substitution at residue 38 of PA (PA-I38M), which was present at the consensus level in nasal swabs from pig #61 at 4 and 5 DPC and is found at sub-consensus levels in other animals (Figure 5). PA-I38M confers resistance to the polymerase inhibitor baloxavir [39]. Another notable consensus-changing nonsynonymous mutation, NA-N307D, was detected in the BALF of #50 at 3 DPC; this mutation may be involved in oseltamivir resistance [39,40]. Several samples also contain viruses harboring HA-D252Y (D236Y in mature H5 numbering and D240Y in H3 numbering); this substitution reached the consensus level in pig #61 oropharyngeal swab at 1 DPC. This residue lies in the HA receptor-binding domain near the 220 loop, but the impact of this particular substitution is not well characterized. One other notable nonsynonymous mutation was HA-S136N (S120N in mature H5 numbering; no directly corresponding amino acid in H3 HA), which was present at 32% frequency in the BALF samples from pig #52 at 3 DPC. This substitution has been shown to increase H5 HA binding to α2,6-linked sialic acids [41]. Also of note, we did not detect emergence of classical mammalian-like mutations in the viral polymerase complex (e.g., PB2-E627K or PB2-D701N) in these pigs, nor did we detect substitutions of other amino acids in HA known to confer effective binding to α2,6-linked sialic acids. Taken together these observations suggest that, although some potentially mammalian-adapting mutations arose in some animals, there is little evidence for extensive additional mammalian adaptation of the bovine-derived HPAI H5N1 B3.13 virus in pigs.

**Figure 5.**
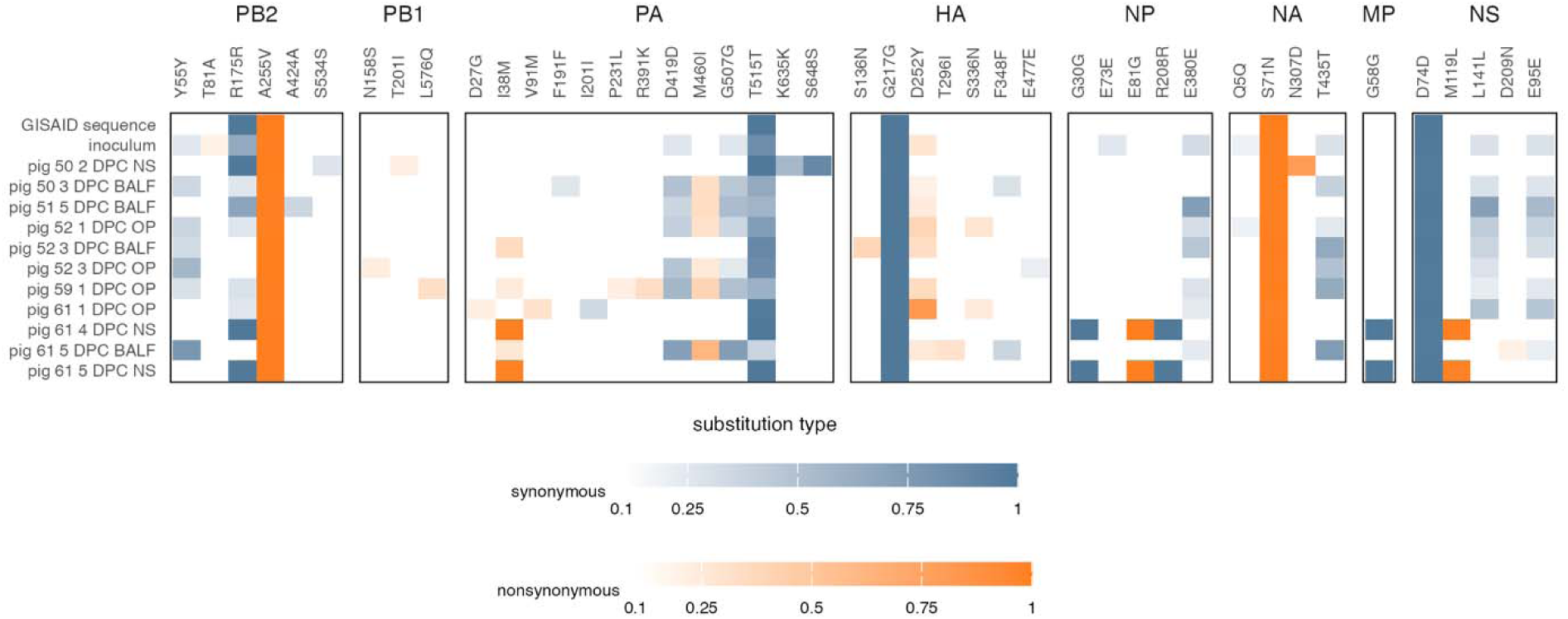
Consensus and subconsensus-level variants across the influenza virus genome in pigs infected with a bovine-derived HPAI H5N1 B3.13 virus. Nonsynonymous and synonymous variants are shown in relation to our analytical pipeline’s standard reference sequence, A/Bovine/Texas/24-029328-01/2024. Amino acid location and identity are labeled. We used the isolate A/Cattle/Texas/063224-24-1/2024 (EPI_ISL_19155861) in this study. The original cattle isolate sequence as deposited in GISAID (GISAID sequence) and the virus stock used to inoculate pigs (inoculum) are also shown in the figure. The inoculum was propagated on bovine uterine epithelial cells for 3 passages at Cornell University and then propagated on MDCK cells for 1 passage at Kansas State University prior to inoculating animals in this study. NS=nasal swab, BALF=bronchoalveolar lavage fluid, and OP=oropharyngeal swab.

## Discussion

IAVs exhibit a variety of host tropisms and can cross species barriers. Since HPAI H5 clade 2.3.4.4b became dominant in wild bird populations, spillovers to mammalian hosts have been frequently reported worldwide, indicating that viruses possessing the 2.3.4.4b HA may have improved ability to replicate in mammals compared to other HPAI lineages [42]. The outbreak of HPAI H5N1 infection in dairy cattle underscores the potential for clade 2.3.4.4b viruses to emerge in mammals. It is critical to understand the potential for cattle-derived viruses to replicate, evolve, and transmit in pigs, a “mixing vessel” species in which novel reassortant viruses with pandemic potential might be generated. Our study demonstrated limited replication of a bovine-derived HPAI H5N1 B3.13 virus in the upper respiratory tracts of pigs. The limited shedding of virus by principal-infected pigs, in turn, led to no transmission to sentinel pigs even after direct contact, although two samples from sentinel pigs were suspect positive in RT-qPCR, which could be due to cross-contamination in the animal room and/or lab. Rather, the bovine-derived HPAI H5N1 B3.13 virus caused self-limiting infection in pigs with virus replication and pathologic alterations mostly affecting the lower respiratory tract. The low virus titers/viral RNA amounts and low viral antigen presence in the lower respiratory tracts demonstrates inefficient virus replication of the bovine-derived HPAI H5N1 B3.13 virus, when compared to the mink-derived H5N1 clade 2.3.4.4b virus or other swine-adapted IAVs [22,43]. Furthermore, detectable antibody responses were identified in only two out of three principal-infected pigs at 14 DPC. Collectively, our results suggest that pigs are only moderately susceptible to the bovine-derived HPAI H5N1 B3.13 virus.

Previous studies have suggested that replication of avian-origin HPAI H5N1 clade 2.3.4.4 viruses are inefficient in the respiratory tracts of pigs [44,45]. In contrast, mammalian isolates of clade 2.3.4.4 exhibit higher respiratory pathogenesis in pigs, when compared to avian-origin clade 2.3.4.4 viruses, which may be due to the presence of mammalian-adapting mutations in the mammalian isolates. For instance, after experimental infection of pigs with A/Mink/Spain/3691-8_22VIR10586-10/2022, an isolate that showed sustained respiratory transmission among farmed mink and possessed a PB2-T271A mutation, efficient viral replication was identified in the lower respiratory tracts, and neutralizing antibodies were detected in 100% of infected pigs within 7 DPC, albeit no transmission to sentinels was observed [22]. Critical mammalian-like adaptations such PB2-E627K and HA-Q222L (mature H5 numbering) were also detected in mink-adapted H5N1-infected pigs, though these remained at low frequencies [22]. In addition, both the A/raccoon/Washington/22-018406-002/2022 H5N1 virus and the A/redfox/Michigan/22-018712-001/2022 H5N1 virus contained the PB2-E627K substitution, and infection of pigs with these viruses led to pig-to-pig transmission, but the transmission efficiency was low: seroconversion was identified in only 1 out of 5 sentinel pigs after co-mingling with 15 principal-infected animals [46]. Although H5N1 viruses isolated from dairy cattle so far mostly lack amino acid substitutions in the polymerase complex previously associated with increased replication in mammalian hosts, such as PB2-E627K and PB2-D701N, they share a different substitution, PB2-M631L, which appears to increase the ability of the polymerase complex to recruit the bovine and human versions of the essential host cofactor ANP32, albeit to a lesser extent than polymerases with PB2-E627K [47]. The PB2-M631L substitution may contribute to the ability of the bovine-derived HPAI H5N1 B3.13 virus to replicate in the respiratory tract of pigs. However, given virus replication kinetics and antibody response, the bovine-derived HPAI H5N1 B3.13 virus was less adapted to the respiratory tract of pigs than the mink-derived HPAI H5N1 virus with the PB2-T271A substitution [22]. Recently, the new genotype D1.1 was also detected in US cattle and characterized as a 4:4 reassortant virus that harbored PB1, HA, M, and NS segments from Eurasian avian lineages and PB2, PA, NP, and NA segments from North American LPAI viruses [9]. Interestingly, the virus retained PB2-D701N mutation that enhances replication in mammalian hosts [48,49], thus the bovine-derived D1.1 virus may present with different replication dynamics in the respiratory tracts of pigs when compared to the bovine-derived B3.13 virus even though the D1.1 virus has not been examined in pigs.

We observed few nonsynonymous mutations in the polymerase complex in cattle-derived H5N1 viruses replicating in swine. The most notable mutation was PA-I38T, which reached consensus in two specimens from one pig (#61) and was detectable at sub-consensus levels in additional specimens from this and other animals. The PA-I38T substitution is known to confer resistance to the antiviral drug baloxavir and has been observed sporadically in other viruses isolated from infected dairy cattle in the current outbreak. Some previous studies suggest that PA-I38T has minimal costs to viral fitness [50], so it is possible that it can arise and persist in some lineages in the absence of baloxavir selection.

In dairy cattle, HPAI H5N1 virus replicates primarily in the mammary tissues and transmission presumably occurs through milk and the milking process. Notably, it is currently understood that bovine mammary tissue mostly expresses “avian-like” α-2,3-linked sialic acid residues [51]. Therefore, serial passages in cattle may not incur selective pressures favoring HA mutations that would increase binding to “mammalian-like” α-2,6-linked sialic acids. In the present study, we observed several mutations that could plausibly alter HA receptor-binding preference of the bovine H5N1 virus. We detected HA-D252Y (D236Y in mature H5 numbering) at the consensus level in one sample from pig #61 and at sub-consensus levels in at least one specimen from every other animal except pig #59. Notably, this substitution was also present at a low level in the inoculum, so it is not clear whether its outgrowth in these animals was due to natural selection or genetic drift. However, this residue is within the receptor-binding domain of HA near to the 220 loop, which is critical for receptor-binding specificity, so it is possible that D252Y represents a mammal-adapting substitution. We also detected HA-S136N (HA-S120N in mature H5 protein numbering), at 32% frequency in a single sample. This mutation was previously shown to increase the binding of H5 HA to α-2,6-linked sialic acids [41], and could therefore have been favored in the pig respiratory tract; although it is notable that this substitution was detected in BALF and not in an upper respiratory tract sample. It is also notable that mutations predicted to confer α-2,6-linked sialic acid binding have been recently detected at sub-consensus levels in two humans with severe infections caused by H5N1 viruses of the D1.1 genotype [52]. It should be noted that in the present study we tested the early bovine isolate from a milk sample that was collected in February 2024. The dairy cattle outbreak remains ongoing as of February 2025, providing additional opportunities for the emergence of mammalian-like mutations. Therefore, it would be valuable to evaluate the pathogenicity and transmissibility of the latest bovine isolates in mammalian hosts in order to compare the level of adaptation to respiratory tracts of mammalian hosts.

In addition to experimental infection of cattle with the bovine-derived HPAI H5N1 B3.13 virus, several groups readily evaluated the pandemic potential of these viruses in well-established animal models. The A/dairy cattle/New Mexico/A240920343-93/2024 strain induced systemic, severe disease in mice after oral ingestion or respiratory infection [53]. Infected ferrets showed clinical signs with systemic viral replication but inefficiently transmitted the virus through the respiratory droplets [53]. Two independent studies supported that the human isolate, A/Texas/37/2024, harboring the PB2-E627K mutation, was more pathogenic and more transmissible than the bovine isolate in ferrets [54,55]. In the primate model, intranasal and intratracheal infection of macaques with A/bovine/OH/B24OSU-342/2024 resulted in mild and severe respiratory disease, respectively, however orogastric inoculation only caused subclinical disease with limited infection and seroconversion [56]. Since the first emergence of the H5N1 B3.13 virus in dairy cattle, the virus has been transmitted from cattle to humans with 41 confirmed human cases and these humans have experienced mild clinical symptoms, such as conjunctivitis, fever and respiratory symptoms with no hospitalization [57]. In contrast, human infections with D1.1 viruses seem to be more severe resulting in the death of one patient [52,58]. Given the severity of clinical outcomes and respiratory tropisms, pigs seem to be better models for the respiratory pathogenesis of HPAI H5N1 clade 2.3.4.4b viruses than mice and ferrets.

In summary, our data suggests that pigs are moderately susceptible to the bovine-derived HPAI H5N1 B3.13 virus but do not transmit to sentinel pigs. Given the important role of pigs in IAV ecology as a mixing vessel for generating the novel reassortant viruses with pandemic potential, enhanced surveillance of pigs is warranted.

## Acknowledgments

We gratefully thank Chester D. McDowell, Daniel W. Madden, Yonghai Li, Shristi Ghimire, Issac Fitz and Mehrnaz Ardalan for assistance with animal and laboratory works. We thank members of the histology laboratories at the Kansas State Veterinary Diagnostic Laboratory and the Louisiana Animal Disease Diagnostic Laboratory for their assistance in histological processing of tissues, slide preparation and staining. We acknowledge BioStone Animal Health for providing the ELISA kit. Funding for this study was provided through grants by USDA NACA 58-3022-3-004, the National Bio and Agro-Defense Facility (NBAF) Transition Fund from the State of Kansas, the USDA Animal Plant Health Inspection Service’s NBAF Scientist Training Program, the AMP and MCB Core of the Center on Emerging and Zoonotic Infectious Diseases (CEZID) of the National Institutes of General Medical Sciences under award number P20GM130448, and the NIAID supported Center of Excellence for Influenza Research and Response (CEIRR) under contract number 75N93021C00016. LSU acknowledges support from an Institutional Development Award (IDeA) from the National Institute of General Medical Sciences of the National Institutes of Health under grant number P20GM130555-5011 (to MC), U.S. Department of Agriculture’s (USDA) National Institute of Food and Agriculture (NIFA) Agriculture and Food Research Initiative and American Rescue Plan Act through USDA Animal and Plant Health Inspection Service (APHIS) competitive grant number 2023-70432-39465 (to MC) and the School of Veterinary Medicine, Louisiana State University (PG009641) (to MC).

## References

[1] Stieneke-Grober A, Vey M, Angliker H, et al. Influenza virus hemagglutinin with multibasic cleavage site is activated by furin, a subtilisin-like endoprotease. EMBO J. 1992 Jul;11(7):2407–14.

[2] World Health Organization. Cumulative number of confirmed human cases for avian influenza A(H5N1) reported to WHO, 2003-2023, 5 January 2023. 2023.

[3] WHO/OIE/FAO H5N1 Evolution Working Group. Toward a unified nomenclature system for highly pathogenic avian influenza virus (H5N1). Emerg Infect Dis. 2008 Jul;14(7):e1.

[4] World Health Organization. Evolution of the influenza A(H5) haemagglutinin: WHO/OIE/FAO H5 Working Group reports a new clade designated 2.3.4.4 2015. Available from: https://www.who.int/publications/m/item/evolution-of-the-influenza-a(h5)-haemagglutinin-who-oie-fao-h5-working-group-reports-a-new-clade-designated-2.3.4.4

[5] Adlhoch C, Fusaro A, Gonzales JL, et al. Avian influenza overview April - June 2023. EFSA J. 2023 Jul;21(7):e08191.

[6] Caserta LC, Frye EA, Butt SL, et al. Spillover of highly pathogenic avian influenza H5N1 virus to dairy cattle. Nature. 2024 Jul 25.

[7] Nguyen T-Q, Hutter C, Markin A, et al. Emergence and interstate spread of highly pathogenic avian influenza A(H5N1) in dairy cattle. bioRxiv. 2024.

[8] Peacock TP, Moncla L, Dudas G, et al. The global H5N1 influenza panzootic in mammals. Nature. 2025 Jan;637(8045):304–313.

[9] Animal and Plant Health Inspection Service. The Occurrence of Another Highly Pathogenic Avian Influenza (HPAI) Spillover from Wild Birds into Dairy Cattle. https://www.aphis.usda.gov/sites/default/files/dairy-cattle-hpai-tech-brief.pdf2025.

[10] Centers for Disease Control and Prevention. HPAI Confirmed Cases in Livestock 2025. Available from: https://www.aphis.usda.gov/livestock-poultry-disease/avian/avian-influenza/hpai-detections/hpai-confirmed-cases-livestock

[11] Centers for Disease Control and Prevention. H5 Bird Flu: Current Situation 2024. Available from: https://www.cdc.gov/bird-flu/situation-summary/index.html#human-cases

[12] Baker AL, Arruda B, Palmer MV, et al. Dairy cows inoculated with highly pathogenic avian influenza virus H5N1. Nature. 2024 Oct 15.

[13] Halwe NJ, Cool K, Breithaupt A, et al. H5N1 clade 2.3.4.4b dynamics in experimentally infected calves and cows. Nature. 2024 Sep 25.

[14] Ma W, Vincent AL, Lager KM, et al. Identification and characterization of a highly virulent triple reassortant H1N1 swine influenza virus in the United States. Virus Genes. 2010 Feb;40(1):28–36.

[15] Nidom CA, Takano R, Yamada S, et al. Influenza A (H5N1) viruses from pigs, Indonesia. Emerg Infect Dis. 2010 Oct;16(10):1515–23.

[16] Choi YK, Nguyen TD, Ozaki H, et al. Studies of H5N1 influenza virus infection of pigs by using viruses isolated in Vietnam and Thailand in 2004. J Virol. 2005 Aug;79(16):10821–5.

[17] Ito T, Couceiro JN, Kelm S, et al. Molecular basis for the generation in pigs of influenza A viruses with pandemic potential. J Virol. 1998 Sep;72(9):7367–73.

[18] Anderson TK, Chang J, Arendsee ZW, et al. Swine Influenza A Viruses and the Tangled Relationship with Humans. Cold Spring Harb Perspect Med. 2021 Mar 1;11(3).

[19] Scholtissek C. Molecular evolution of influenza viruses. Virus Genes. 1995;11(2-3):209–15.

[20] Mena I, Nelson MI, Quezada-Monroy F, et al. Origins of the 2009 H1N1 influenza pandemic in swine in Mexico. Elife. 2016 Jun 28;5.

[21] Animal and Plant Health Inspection Service. Federal and State Veterinary Agencies Share Update on HPAI Detections in Oregon Backyard Farm, Including First H5N1 Detections in Swine. https://www.aphis.usda.gov/news/agency-announcements/federal-state-veterinary-agencies-share-update-hpai-detections-oregon2024.

[22] Kwon T, Trujillo JD, Carossino M, et al. Pigs are highly susceptible to but do not transmit mink-derived highly pathogenic avian influenza virus H5N1 clade 2.3.4.4b. Emerg Microbes Infect. 2024 Dec;13(1):2353292.

[23] Sponseller BA, Strait E, Jergens A, et al. Influenza A pandemic (H1N1) 2009 virus infection in domestic cat. Emerg Infect Dis. 2010 Mar;16(3):534–7.

[24] Gauger PC, Vincent AL. Serum Virus Neutralization Assay for Detection and Quantitation of Serum Neutralizing Antibodies to Influenza A Virus in Swine. Methods Mol Biol. 2020;2123:321–333.

[25] Lail AJ, Vuyk WC, Machkovech H, et al. Pasteurized retail dairy enables genomic surveillance of H5N1 avian influenza virus in United States cattle. medRxiv. 2024.

[26] Quick J, Grubaugh ND, Pullan ST, et al. Multiplex PCR method for MinION and Illumina sequencing of Zika and other virus genomes directly from clinical samples. Nat Protoc. 2017 Jun;12(6):1261–1276.

[27] Di Tommaso P, Chatzou M, Floden EW, et al. Nextflow enables reproducible computational workflows. Nat Biotechnol. 2017 Apr 11;35(4):316–319.

[28] Martin M. Cutadapt removes adapter sequences from high-throughput sequencing reads. EMBnetjournal. 2011;17(1).

[29] Li H. Minimap2: pairwise alignment for nucleotide sequences. Bioinformatics. 2018 Sep 15;34(18):3094–3100.

[30] Burrough ER, Magstadt DR, Petersen B, et al. Highly Pathogenic Avian Influenza A(H5N1) Clade 2.3.4.4b Virus Infection in Domestic Dairy Cattle and Cats, United States, 2024. Emerg Infect Dis. 2024 Jul;30(7):1335–1343.

[31] Castellano S, Cestari F, Faglioni G, et al. iVar, an Interpretation-Oriented Tool to Manage the Update and Revision of Variant Annotation and Classification. Genes (Basel). 2021 Mar 8;12(3).

[32] Cingolani P, Platts A, Wang le L, et al. A program for annotating and predicting the effects of single nucleotide polymorphisms, SnpEff: SNPs in the genome of Drosophila melanogaster strain w1118; iso-2; iso-3. Fly (Austin). 2012 Apr-Jun;6(2):80–92.

[33] Danecek P, Bonfield JK, Liddle J, et al. Twelve years of SAMtools and BCFtools. Gigascience. 2021 Feb 16;10(2).

[34] Shen W, Sipos B, Zhao L. SeqKit2: A Swiss army knife for sequence and alignment processing. Imeta. 2024 Jun;3(3):e191.

[35] Hall M. Rasusa: Randomly subsample sequencing reads to a specified coverage. Journal of Open Source Software. 2022;7(69).

[36] Rognes T, Flouri T, Nichols B, et al. VSEARCH: a versatile open source tool for metagenomics. PeerJ. 2016;4:e2584.

[37] R Core Team. R: A Language and Environment for Statistical Computing. R Foundation for Statistical Computing. https://www.R-project.org/. 2024.

[38] Grubaugh ND, Gangavarapu K, Quick J, et al. An amplicon-based sequencing framework for accurately measuring intrahost virus diversity using PrimalSeq and iVar. Genome Biol. 2019 Jan 8;20(1):8.

[39] Nguyen HT, Chesnokov A, De La Cruz J, et al. Antiviral susceptibility of clade 2.3.4.4b highly pathogenic avian influenza A(H5N1) viruses isolated from birds and mammals in the United States, 2022. Antiviral Res. 2023 Sep;217:105679.

[40] Stoner TD, Krauss S, DuBois RM, et al. Antiviral susceptibility of avian and swine influenza virus of the N1 neuraminidase subtype. J Virol. 2010 Oct;84(19):9800–9.

[41] Wang W, Lu B, Zhou H, et al. Glycosylation at 158N of the hemagglutinin protein and receptor binding specificity synergistically affect the antigenicity and immunogenicity of a live attenuated H5N1 A/Vietnam/1203/2004 vaccine virus in ferrets. J Virol. 2010 Jul;84(13):6570–7.

[42] World Health Organization. Assessment of risk associated with recent influenza A(H5N1) clade 2.3.4.4b viruses. 2022.

[43] Kwon T, Artiaga BL, McDowell CD, et al. Gene editing of pigs to control influenza A virus infections. Emerg Microbes Infect. 2024 Jul 31:2387449.

[44] Kaplan BS, Torchetti MK, Lager KM, et al. Absence of clinical disease and contact transmission of HPAI H5NX clade 2.3.4.4 from North America in experimentally infected pigs. Influenza Other Respir Viruses. 2017 Sep;11(5):464–470.

[45] Graaf A, Piesche R, Sehl-Ewert J, et al. Low Susceptibility of Pigs against Experimental Infection with HPAI Virus H5N1 Clade 2.3.4.4b. Emerg Infect Dis. 2023 Jul;29(7):1492–1495.

[46] Arruda B, Baker ALV, Buckley A, et al. Divergent Pathogenesis and Transmission of Highly Pathogenic Avian Influenza A(H5N1) in Swine. Emerg Infect Dis. 2024 Apr;30(4):738–751.

[47] Dholakia V, Quantrill JL, Richardson S, et al. Polymerase mutations underlie early adaptation of H5N1 influenza virus to dairy cattle and other mammals. bioRxiv. 2025.

[48] Li Z, Chen H, Jiao P, et al. Molecular basis of replication of duck H5N1 influenza viruses in a mammalian mouse model. J Virol. 2005 Sep;79(18):12058–64.

[49] Gao Y, Zhang Y, Shinya K, et al. Identification of amino acids in HA and PB2 critical for the transmission of H5N1 avian influenza viruses in a mammalian host. PLoS Pathog. 2009 Dec;5(12):e1000709.

[50] Imai M, Yamashita M, Sakai-Tagawa Y, et al. Influenza A variants with reduced susceptibility to baloxavir isolated from Japanese patients are fit and transmit through respiratory droplets. Nat Microbiol. 2020 Jan;5(1):27–33.

[51] Kristensen C, Jensen HE, Trebbien R, et al. Avian and Human Influenza A Virus Receptors in Bovine Mammary Gland. Emerg Infect Dis. 2024 Sep;30(9):1907–1911.

[52] Jassem AN, Roberts A, Tyson J, et al. Critical Illness in an Adolescent with Influenza A(H5N1) Virus Infection. N Engl J Med. 2024 Dec 31.

[53] Eisfeld AJ, Biswas A, Guan LZ, et al. Pathogenicity and transmissibility of bovine H5N1 influenza virus. Nature. 2024 Sep 12;633(8029).

[54] Gu C, Maemura T, Guan L, et al. A human isolate of bovine H5N1 is transmissible and lethal in animal models. Nature. 2024 Oct 28.

[55] Pulit-Penaloza JA, Belser JA, Brock N, et al. Transmission of a human isolate of clade 2.3.4.4b A(H5N1) virus in ferrets. Nature. 2024 Oct 28.

[56] Rosenke K, Giffin A, Kaiser F, et al. Pathogenesis of bovine H5N1 clade 2.3.4.4b infection in Macaques. Nature. 2025 Jan 15.

[57] Garg S, Reinhart K, Couture A, et al. Highly Pathogenic Avian Influenza A(H5N1) Virus Infections in Humans. N Engl J Med. 2024 Dec 31.

[58] Centers for Disease Control and Prevention. CDC Confirms First Severe Case of H5N1 Bird Flu in the United States. https://www.cdc.gov/media/releases/2024/m1218-h5n1-flu.html2024.

