## Supplemental Material for "Pathogenicity and transmissibility of bovine-derived HPAI H5N1 B3.13 virus in pigs"

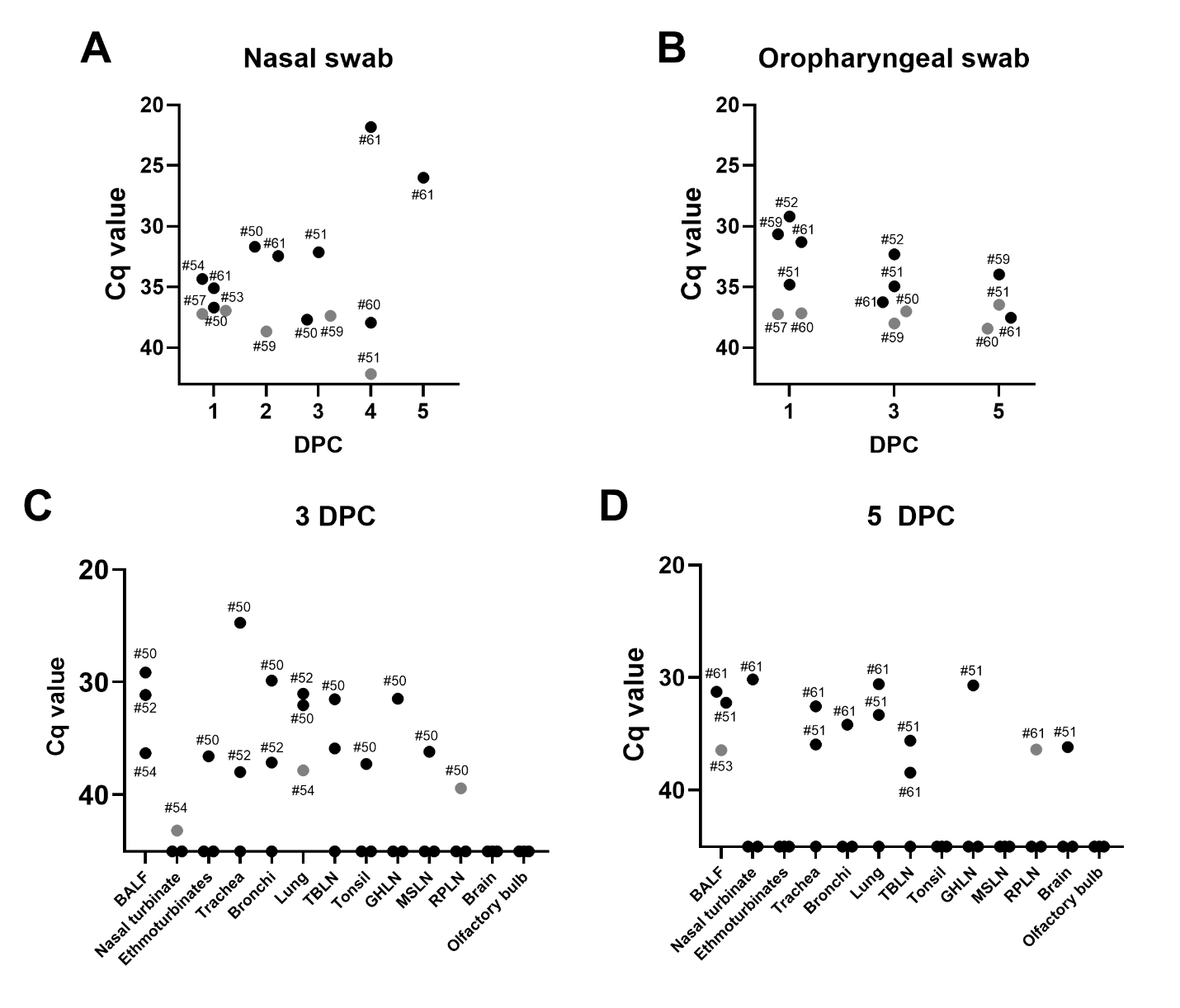

**Supplementary figure 1**. Viral RNA detection in (A) nasal swab, (B) oropharyngeal swab, and tissues at 3 DPC (C) and 5 DPC (D) from pigs infected with a bovine-derived HPAI H5N1 B3.13 virus. Only positive (closed) and suspect-positive (grey) samples of principal pigs out of n=9 at 1 to 3 DPC and n=6 at 4 and 5 DPC were shown and labeled with pig ID (A and B). Tissue samples from three principal pigs at 3 DPC and 5 DPC were tested, and positive and suspect-positive samples are represented as closed and grey circles, respectively. Formalin-fixed trachea and bronchi collected from #50 at 3 DPC were tested for RT-qPCR. TBLN=tracheobronchial lymph node, GHLN=gastrohepatic lymph node, MSLN=mesenteric lymph node, RPLN=retropharyngeal LN.

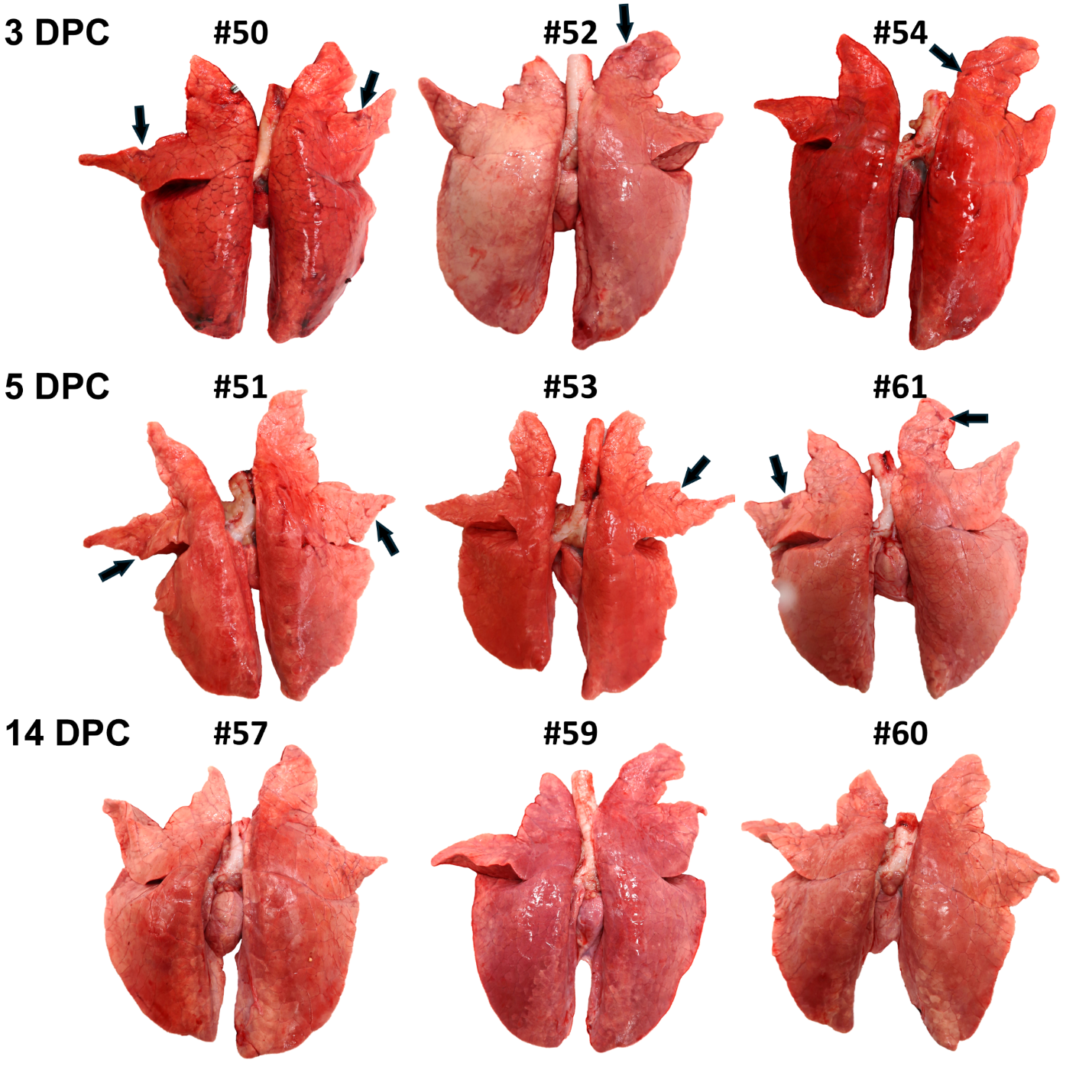

**Supplementary figure 2.** Gross changes in the lungs of a bovine-derived HPAI H5N1 B3.13 virus-infected pigs at 3, 5 and 14 DPC. The arrows represent macroscopic lung lesions showing sporadic dark red and depressed lobules representing regions of lobular atelectasis that are seen in IAV-infected pigs.

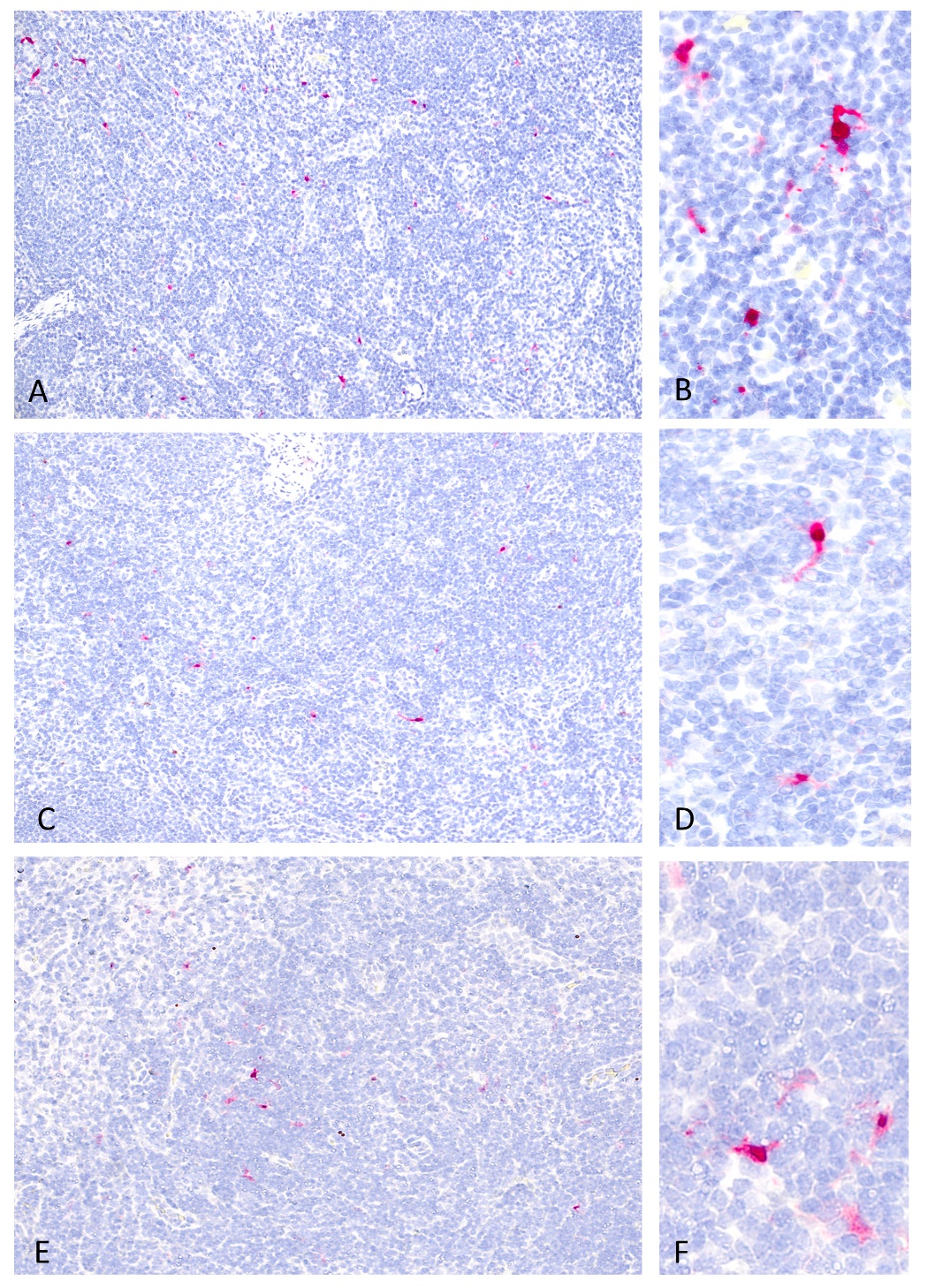

**Supplementary figure 3.** Influenza A virus-specific immunohistochemistry for detecting H5N1 antigens. Positive signals were detected in large pleomorphic cells within the tracheobronchial lymph nodes at 3 (A and B), 5 (C and D) and 14 DPC (E and F). Influenza A virus NP antigen was less abundant but had a similar cellular distribution at 14 DPC compared to 3 and 5 DPC

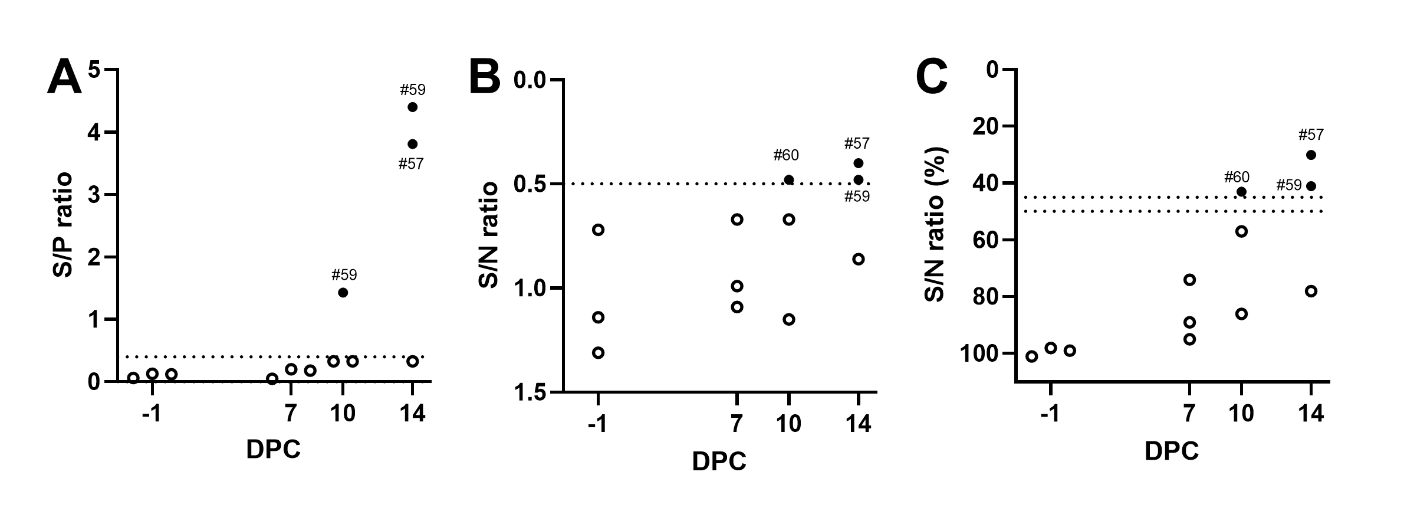

**Supplementary figure 4**. Antibody responses determined by ELISA in pigs infected with a bovine-derived HPAI H5N1 B3.13 virus. Commerical ELISA was performed according to the manufacturers’ instructions: (A) ID Screen® Influenza A Nucleoprotein Swine Indirect, Innovative Diagnostics, France, (B) IDEXX Swine Influenza Virus Ab Test, IDEXX, USA and (C) ID Screen® Influenza A Antibody Competition Multi-Species, Innovative Diagnostics, France. The positive samples were shown as closed circles with pig ID: open circles represent negative. The dash lines represent the cut-off values of each ELISA: (A) >0.4 of the S/P ratio, (B) <0.5 of the S/N ratio, and (C) <45%: positive, 45-50%: doubtful, and >50%: negative.

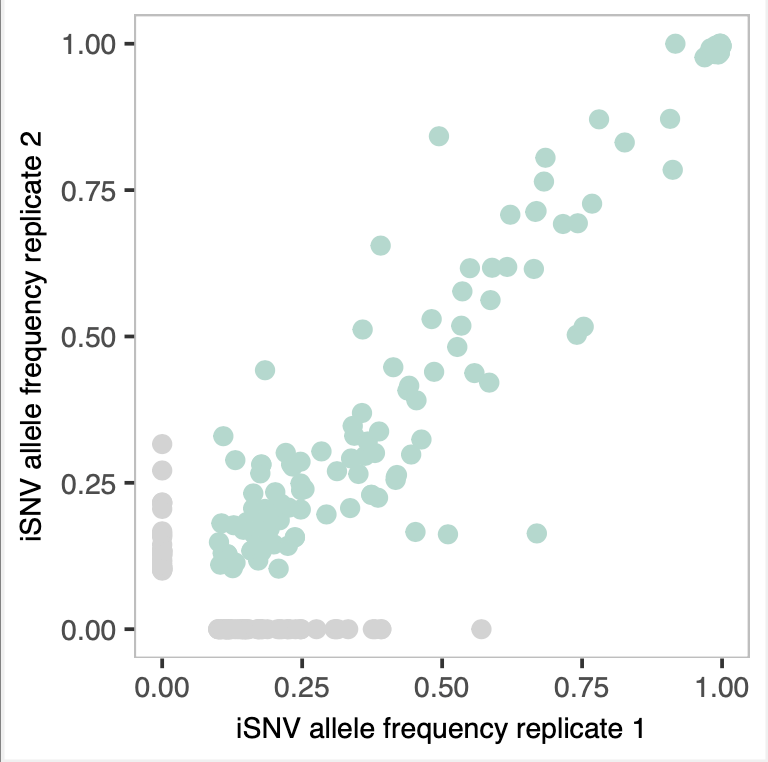

**Supplementary figure 5. Pairwise comparison of iSNV frequencies in technical replicates.** iSNV frequencies in replicate 1 are shown on the x-axis and frequencies in replicate 2 are shown on y-axis. Gray colored iSNVs are present in only 1 replicate, and were excluded from subsequent analyses.
